# Profiling co-occurrent morphological phenotypes and their degree of expression severity in vacuolated cells by holo-tomographic flow cytometry and fractal analysis

**DOI:** 10.64898/2026.02.09.704871

**Authors:** Marika Valentino, Giusy Giugliano, Daniele Pirone, Fabrizio Licitra, Fulvia Vitale, Pasquale Memmolo, Lisa Miccio, Massimo D′Agostino, Pietro Ferraro, Vittorio Bianco

## Abstract

Cells are complex systems characterized by large phenotype heterogeneity. Conventional single cell classification approaches usually separate cells expressing a certain phenotype (e.g. associated to a disease condition) from the healthy control. However, multiple phenotypes typically coexist within the same cell as a result of complex intracellular interactions, machineries, functioning and external stimuli. Here we use label-free optical microscopy, powered by AI, to investigate how morphological phenotypes co-occur within vacuolated cells. Cytoplasmic vacuoles are important hallmarks of several pathological states (e.g. lysosomal storage diseases, viral infections, cancer). We rely on Holo-Tomographic Flow Cytometry (HTFC) to obtain 3D refractive index tomograms of vacuolated cells in continuous flow. Then, we propose a strategy to reduce the dimensionality of the tomogram using cross-sectioning and minimum intensity projection maps. We extract a set of morphological, refractive index-based, and fractal parameters demonstrating that the complex heterogeneity of vacuole patterns can be captured and can foster classification based on interpretable features. For training the AI, biologist domain-experts provided annotation of the different morphological phenotypes expressed and ranked them in terms of expression severity from the tomographic observations. Thus, we introduce a pipeline for morphometric phenotype profiling, in which each cell is associated with a 7-digits classification code representing the combination of coexisting phenotypes it expresses and their expression severity levels.

## 1. Introduction

Either from the surface or from the intracellular perspective, within the same or between different cell populations, substantial variation can be observed due to high cellular heterogeneity [1]. A specific combination of these features often serves as a unique fingerprint, allowing biologists and pathologists to classify cell types, determine their cell-cycle stage, or identify abnormal states [2]. Fluorescence microscopy methods coupled to conventional machine learning or deep learning analysis can be used to identify phenotypes within a cell or a cell line [3-6]. However, these marker-based approaches can be invasive and alter the natural functioning of the cells under test, and thus the phenotypes to be identified, while lab protocols and operator dependencies still exist [7]. By using variations in the refractive index (RI) inside the specimen as an endogenous contrast mechanism, Digital Holography in transmission microscopy configuration provides the 2D morphology and quantitative Phase-Contrast Map (PCM) of label-free samples [8-10]. However, an integral 2D PCM cannot map in full the 3D morphology of a cell, its organelles and the heterogeneity of its morphological phenotypes. Holo-Tomographic Flow Cytometry (HTFC) combines the PCM corresponding to multiple orientations of the same cell while it flows and rotates in controlled manner inside a microfluidic channel to provide the 3D Refractive Index (RI) tomogram [11-14]. Combining microfluidics and holographic tomography (HT) has a threefold advantage over conventional static approaches [15-17]: i) it accesses high-throughput in flow cytometry mode; ii) cells that naturally live in suspension can be imaged in suspension without altering their morphology, and thus the 3D positioning of their organelles can be mapped. iii) full rotation sequences are obtainable, thus quasi-isotropic resolution can be accessed due to the lack of severe missing cone problems as for 3D static HT. Recently, HTFC has been demonstrated to provide tomograms with one more channel of information, i.e. label-free intracellular specificity for multiple cell sub-compartments including nuclei, cytoplasmic vacuoles, lysosomes, lipid droplets, peri-vacuolar and peri-nuclear regions, cytoplasm [18-22]. This type of 3D segmentation is based on a statistical analysis of the tomogram voxels and can be obtained from the 3D data only. Accurately identifying the nucleus region is of interest since its size and shape are important biomarkers (e.g. an abnormal nucleus/cytoplasm biovolume ratio is a hallmark for cancer cells) [23]. While normal cells maintain nuclear size within a defined range, altered nuclear size and shape are associated with a variety of diseases. Cancer-associated changes in nuclear morphology may disrupt normal chromatin positioning, gene expression, and DNA damage pathways, potentially contributing to disease progression. Moreover, mutations in nuclear proteins contribute to other diseases, including muscular dystrophy, premature ageing, laminopathies, amyotrophic lateral sclerosis and frontotemporal dementia [24]. The presence in mammalian cells of cytoplasmic vacuoles is of particular interest as well, since these sub-structures are frequently linked with pathogenic states [25]. Numerous illnesses, such as lysosomal storage disorders (LSDs) [26], cancer [27,28], and viral infections [29,30], have been associated to their irregular accumulation. Therapeutic drugs can accumulate in lysosomes during specific therapies, causing reduced effectiveness and the enlargement of the lysosomal compartment, thereby forming cytoplasmic vacuoles. This impairs the function of the lysosomes and increases resistance to therapy [31,32]. Thus, the identification of vacuolar phenotypes in lysosomes and the cytoplasm is diagnostically important, which in turn highlights the need for cytometry methods that can quantify the vacuolization degree [33]. All the above-mentioned cell phenotypes manifest with morphological alterations and can be more or less expressed in the same cell population, which leads to a large heterogeneity *per se*. Above all, they can coexist within the same cell, each with a certain expression severity. As a result, the classical “healthy vs. sick” cell classification is inappropriate to describe this intra- and inter-cellular heterogeneity of morphological phenotypes.

In this study, we exploit HTFC with nucleus specificity capability and machine learning to map the sequence of morphological phenotypes coexisting in each single cell within a population under test. At this scope, we first apply the Computational Segmentation based on Statistical Inference (CSSI) algorithm [18-22] to identify the nucleus region from the 3D RI tomogram. Then, we introduce a method to reduce the dimensionality of the tomogram, with the scope to handle efficiently 3D data without losing insightful information. We exploit a combination of cross-sectional and Minimum Intensity Projection (MIP) maps for the scope. We use these reduced-dimensionality tomographic data and the nucleus information gathered at the previous step to extract a set of interpretable morphometric, refractive index-based, and fractal features [34]. Thus, we use them to classify the morphological phenotypes and their expression severity within each cell. The combination of morphological traits expressed at various levels in a cell is a sequence that maps the unique fingerprint of the line it belongs, its status, complex machineries, and all in all its phenotype. Figure 1 summarizes the proposed analysis pipeline. As a proof of concept of the capabilities of AI-aided HTFC, we investigate the morphometric signatures associated with the process of vacuolization in U937 monocyte cells. The choice of vacuolated cells for this analysis is mostly driven by the importance of this process and its association to widespread and severe pathologies, as discussed above. Besides, we are able to control and guide the vacuolization process in monocyte cells [35,36] in order to obtain enough examples of this set of phenotype sequences with variable severity and access a reference ground truth. We exposed cells to increasing concentrations of vacuoline, a compound that induces vacuole formation. Expert biologists inspected the full tomographic dataset. They indicated for each cell a sequence of the major morphological phenotypes expressed and ranked them according to four levels of severity (i.e. Level=0 means absence of a phenotype, Level=3 means phenotype very severely expressed). The phenotypes considered in the proposed analysis (herein named as A-G) are the (A) altered nuclear morphology, (B) evaluation of the arise of cytoplasmic vacuoles in terms of size and number, (C) irregular spatial distribution of the vacuoles in the cell volume, (D) Vacuole area relative to cell volume, (E) Cytosol-to-nucleus ratio as a measure of nuclear/cytoplasmic balance, (F) Cell size, (G) Cell ovality/eccentricity.

**Figure 1.**
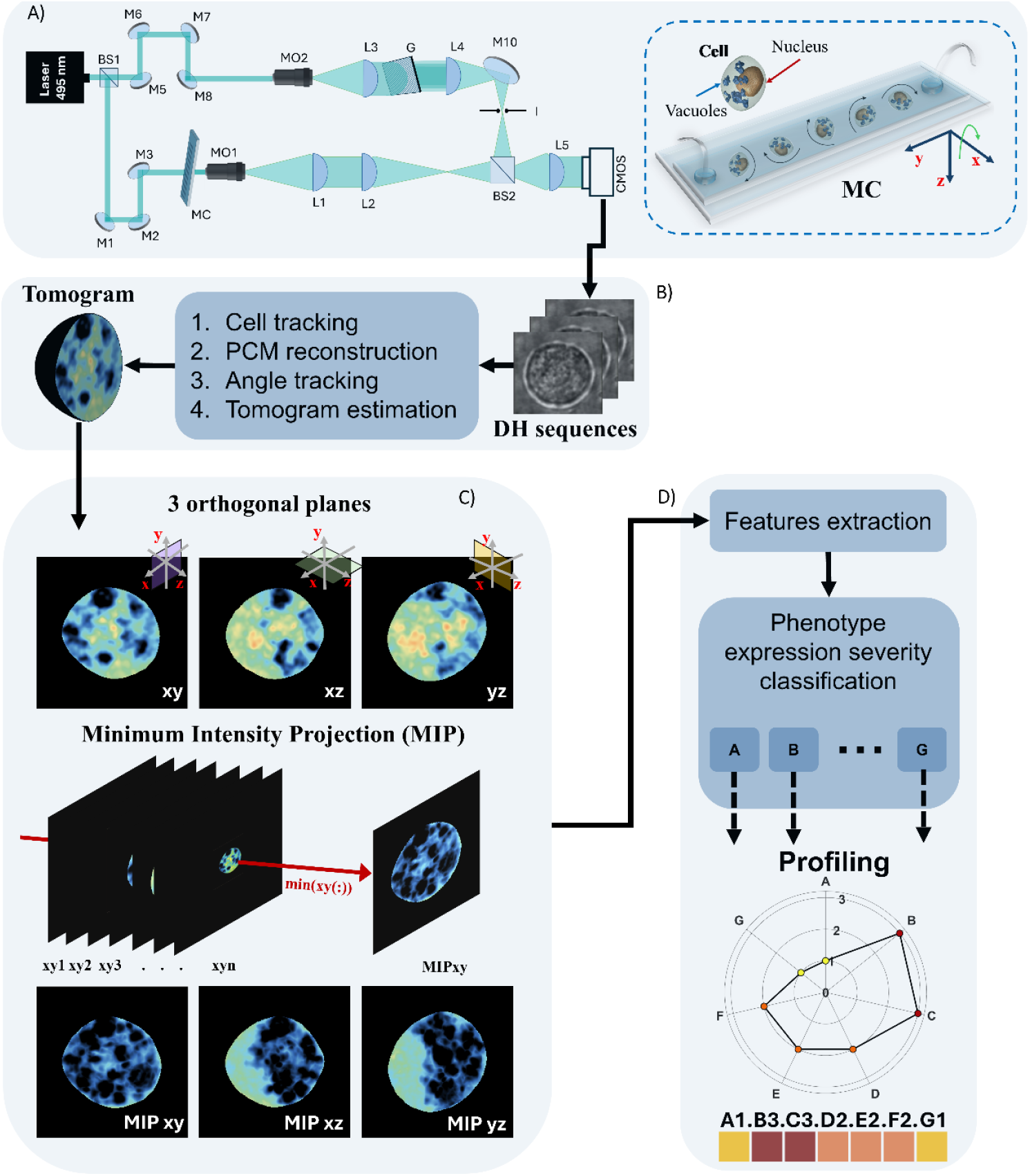
Working pipeline. As cells go through the microfluidic channel (MC), they undergo natural rotation along the x-axis, which provides multiple angular views recorded as holographic videos using the HTCF optical setup. A) Sketch of the experimental arrangement: BS, beam splitters; ND, neutral density filter; M, mirrors; MO, microscope objectives; L, achromatic doublets; G, diffraction grating; I, iris diaphragm. B) The hologram sequences are converted into PCMs and subsequently processed to reconstruct 3D RI tomograms. C) Dimensionality reduction: from each tomogram, cross-sections along the three orthogonal planes (xy, xz, yz) are extracted, along with Minimum Intensity Projection (MIP) maps. D) Profiling: morphological parameters, RI-based measurements, and fractal descriptors are derived from the dimensionality-reduced auxiliary maps. The resulting feature set allows profiling the morphological phenotypes coexisting in each single cell and the assessment of their expression severity levels. The compresence of these phenotypes is a unique fingerprint of the cell, here summarized with a 7 digits sequence and a spider-graph for each cell.

A more detailed description of the diagnostic relevance of the selected phenotypes and their link to dysfunctions and pathological conditions is reported in the Supplementary information. Of course, some of them can be absent/not identified in some cell depending on the treatment it has undergone. It is worth noticing that this set of morphological phenotypes is observable thanks to the combined information content provided by the 3D tomographic cell characterization and the nuclear 3D shape estimation. We have observed that vacuoline treatment leads to heterogeneous phenotypic responses and varying degrees of vacuolization, well mimicking the variability occurring in the case of vacuolization-associated pathologies. We use this extensive dataset labelling to train and validate our classification models. The result is a morphological phenotype profiling, i.e. each cell passing through the HTFC system is mapped with a 7-digits code resulting from the combination of morphological phenotypes it expresses, and the expression severity levels stemming from their compresence.

## 2. Methodology

### 2.1 Outline of HTFC

In our HTFC setup (Fig. 1 A), cells suspended in continuous flow pass through the MC while illuminated by a coherent laser beam in off-axis holographic microscopy configuration (see Supplementary Information) [11,19]. During their transit, cells undergo rotation. This rotational motion allows the acquisition of sequences of digital holograms at multiple viewing angles without changing the illumination direction in the setup (MC sketched in Fig. 1 A). Each hologram encodes the optical phase delay generated by the RI distribution of the cell. Assuming steady flow along the y-axis, ensured by the syringe pump, cells rotate around the x-axis. Once a target cell is tracked, a sequence of square sub-holograms is cropped from consecutive frames, each centered on the cell during its rotation. These cropped holograms serve as input for the phase-reconstruction algorithm [11], which includes focus distance estimation (Tamura coefficient optimization), aberration correction, noise suppression, and phase unwrapping (Fig. 1 B) [11,37,38]. The output is a series of PCMs associated with successive rotation angles. Angle assignment begins from an arbitrary 0° reference for the first PCM and proceeds according to the estimation algorithm described in [38], thereby associating each phase map with a specific orientation angle (Fig. 1 B). By combining the set of PCMs and using the estimated angle sequence, tomographic inversion algorithms such as the filtered back-projection [39,13,15], return a volumetric map of the cell RI distribution. Figure 1 B summarizes the main processing steps that allow passing from the holographic sequence to the cell RI tomogram. The experimental setup is detailed in the Supplementary Information (Fig. S1). Moreover, we enrich this 3D information content by adding nucleus specificity to the 3D RI tomogram of each cell, using the CSSI algorithm [18-22]. This approach allowed us segmenting the nucleus voxels from the 3D RI tomograms.

### 2.2 Sample preparation

U937 cells from ATCC biobank (ATCC-CRL-1593.2) were routinely grown at 37°C, 5% CO2, in RPMI-1640 medium supplemented with 10% fetal bovine serum (FBS; Sigma-Aldrich), 100 U/ml Penicillin/Streptomycin, and 2 mM L-Glutamine (L-Gln; Sigma-Aldrich). Cells were treated with Vacuolin-1 (MedChemExpress) starting from the storage concentration of 10µM dissolved in DMSO. After treatment, cells were diluted to 10^5^ cells/ml and analysed by HTFC.

### 2.3 Dimensionality reduction

In order to efficiently handle the 3D content of a tomogram, here we propose the use of a set of 2D auxiliary maps obtained from the tomogram to extract the features exploitable for the classification of morphological phenotypes (Fig. 1 C). In principle, any tomogram cross-section could be used to extract features, and the full information content would be obtained only considering the full 3D stack. Moreover, it is worth pointing out that accessing in the first place the 3D tomogram is pivotal for the proposed analysis, since the nucleus and in turn the vacuolar regions are identifiable in marker-free mode only from the tomogram by employing the CSSI method [18-22]. However, we operate a dimensionality reduction for extracting the features to streamline the analysis pipeline and propose a robust solution that scales up well with the increase of the cell throughput. Dimensionality reduction facilitates the extraction of multi-scale fractal parameters that are very descriptive of morphometric phenotypes [34,40-42] but require long computational times when applied to 3D stacks. In particular, we use the RI slices corresponding to the three orthogonal cross-sectional planes (*xy, xz*, and *yz*) of each 3D RI tomogram, namely *S*_*xy*_, *S*_*xz*_, *S*_*yz*_, along with the Minimum Intensity Projection (MIP) maps calculated across the three main axis. Features are then extracted from the stack of 2D matrixes Σ = {*S*_*xy*_, *S*_*xz*_, *S*_*yz*_, *MIP*_*xy*_, *MIP*_*xz*_, *MIP*_*yz*_}. This choice lowers the risk of overfitting and decreases the computational cost of downstream tasks as mentioned above. For instance, computing the 3D lacunarity takes approximately 11s per cell using a desktop computer with 64GB RAM and Intel-i9 processor, while this time lowers to 0.04s per cell if the 2D lacunarity has to be calculated. Considering the use of the stack of matrixes Σ, the computational time for the lacunarity calculation is 0.24s, i.e. it takes only the 2.18% of the time and scales up much better with the increase of the cell number. By limiting also the projections to the three orthogonal planes, we balanced computational efficiency with the preservation of biologically meaningful information.

### 2.4 Vacuoles and nuclei segmentation

After obtaining the RI slices (an example is reported in Fig. 2 A-C) and the corresponding MIP maps along the three principal planes (Fig. 1 C), vacuoles were segmented individually in each plane for each cell, as shown in Fig. 2 A1-C1. Segmentation was performed automatically by using a global thresholding approach. Specifically, the threshold was set to the median of the 20th percentile of the RI values computed across all analysed cells. This fixed threshold was then applied to all RI and MIP maps, ensuring uniformity in the detection process across different imaging modalities and experimental conditions.

**Figure 2.**
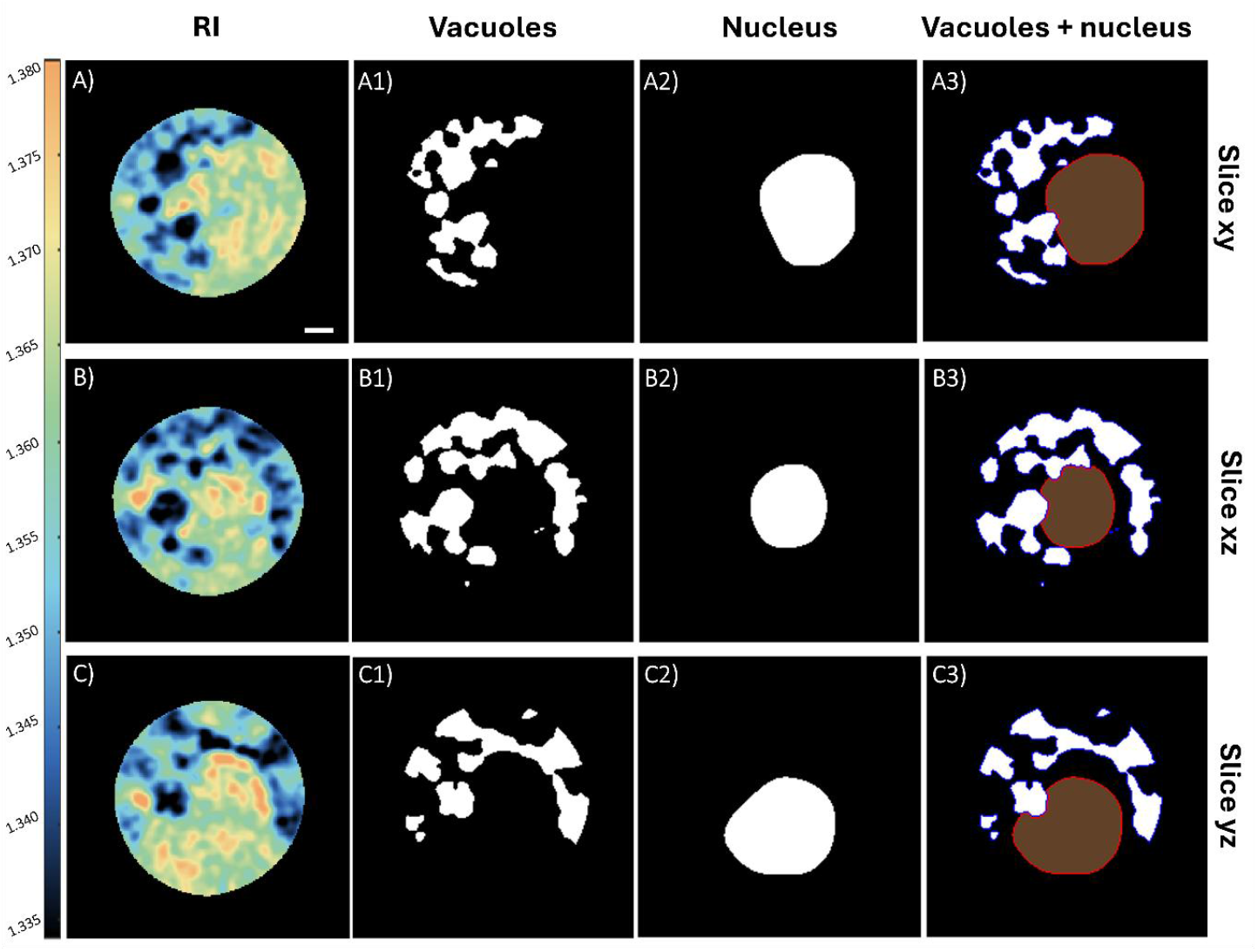
Segmentation of vacuoles and nucleus from RI and MIP maps. (A–C) Orthogonal RI slices of a representative cell. (A1–C1) Vacuole segmentation in each plane, obtained through an automated global thresholding strategy. (A2–C2) Nuclear segmentation performed with the CSSI algorithm [M33], shown in the three main cross-sections. (A3–C3) Fusion of vacuole and nuclear masks using a logical OR operation, generating irregular, non-convex nuclear contour that reflect local membrane deformation due to vacuole proximity. The colour-bar represents the RI values.

After applying the CSSI algorithm [18-22], a 3D binary mask of the cell nucleus was obtained. As in the previous analyses, the three principal planes (*xy, xz* and *yz*) of this segmented volume were considered to obtain the corresponding nuclear cross-sections, Fig. 2 A2-C2. Noteworthy, cytoplasmic vacuoles were found to be in close proximity to the nuclear membrane, often appearing to compress and deform the nucleus by exerting local pressure on its surface. To account for this phenomenon, the binary masks of the nucleus were combined with the vacuoles masks using a logical OR operation (Fig. 2 A3-C3). This fusion resulted in irregular, non-convex nuclear masks that more accurately reflect the membrane deformation caused by adjacent vacuoles along the three orthogonal planes.

### 2.5 Features extraction and fractal analysis

For each cell, the dataset was built by extracting features from: *(i)* the three orthogonal slices belonging to the RI tomogram; *(ii)* the three orthogonal slices belonging to the corresponding vacuoles segmentation maps; *(iii)* the three orthogonal slices belonging to nucleus volume; *(iv)* the three orthogonal slices belonging to the MIP maps; *(v)* the three orthogonal slices belonging to the corresponding vacuoles segmentation maps, thus totalizing 15 auxiliary images for cell (feature extraction step, Fig. 1 D). Although these three orthogonal slices belong to the same tomographic volume, we treated them as independent samples. This approach acts as a form of data augmentation, providing different morphological perspectives for learning and improving model generalization. For each orthogonal plane, we extracted a total of 37 features, including morphological, RI-based, and fractal descriptors. We explored features derived from fractal geometry since this set of descriptors decipher well the inner complexity of the auxiliary maps. In particular, vacuole segmentation maps present irregular patterns and can be interpreted as cell segmentation masks with internal holes. Fractal analysis of this binary pattern, can capture geometry complexity, spatial inhomogeneity, and structural fragmentation at multiple scales [34,40-42]. Among the fractal features considered, we included the fractal dimension, the lacunarity, and the vertex density [42]. These descriptors, in our findings, provide complementary information about the contour complexity, statistical distribution of the RI holes and structural irregularity of the vacuole patterns. The definition of each feature is reported in Table S1 of the Supplementary Information.

After feature extraction, a supervised feature selection strategy was applied to reduce redundancy and retain the most informative features for the classification tasks. Specifically, we used the Minimum Redundancy Maximum Relevance (mRMR) algorithm [43] to rank features based on their relevance to the target classes and their redundancy with respect to each other. Features with a selection score above the 75th percentile of the total distribution were kept. This percentile-based thresholding ensured an automatic and adaptive selection process, keeping only the most significant and non-redundant descriptors for further analysis.

### 2.6 Classification approach

To perform degree of expression severity classifications for each phenotype, we designed a dataset and modelling pipeline devoted to solve two distinct tasks: binary classification and multiclass classification. For each task, we first performed classification using the entire features set, then classification was repeated on a features subgroup after features selection.

Instead of classifying cells as in conventional approaches, here we attempt classification separately for each of the seven morphological phenotypes considered (phenotype expression severity classification, Fig. 1 D). Expression severity of each phenotype was grouped in four levels (i.e. 0, 1, 2, 3), where the 0 level means that a phenotype is not present/expressed in the single cell. Details on the process of labelling of the coexisting morphological phenotypes in the training set by domain-experts are provided in the Supplementary Information. In the binary classification task, severity levels were grouped into two main classes: [0, 1] as class 0 (weakly or not expressed), and [2, 3] as class 1 (high expression severity). In the multiclass identification task, all four severity levels were treated as distinct classes, resulting in a four-class problem. For each classification problem, individual classifiers were trained and tested by using the Matlab® Classification Learner App [44]. The best-performing model for each case was based on the highest test accuracy. Finally, to improve predictions at the cell level, a Majority Voting (MV)) strategy [45] was applied across the three slices belonging to the same cell. The final phenotype severity prediction for each test cell was assigned based on the most frequent class predicted among its three slices. For example, if the cell “number 345” is under test and the phenotype C is classified as severity level 0 in two out of three slices, the profile code of the “cell 345” is constructed by adding C0 to its morphological sequence. For each cell, classification of all morphological phenotypes is performed as described, which yields the full cell morphological profiling.

## 3. Experimental results

### 3.1 Dataset description

A total of 618 cells was collected for this study, including wild-type (WT) cells and cells treated with vacuoline at concentrations of 0.1 μM, 1 μM, and 5 μM, of which the latter were observed at two time points: 1 hour and 2.5 hours post-treatment. For each cell, the three principal tomographic planes were considered and the resulting 1 × 37 feature vectors were treated as independent samples, resulting in a total dataset of 1854 samples (as listed in Table S1 of the Supplementary Information). The dataset was then split into 80% for training and 20% for testing. The training set included the 10-fold cross-validation. The test set comprised 372 feature vectors, corresponding to 124 cells (each represented by the vectors of the three main orthogonal slices), allowing classification performance to be evaluated both at the feature vector level and at the morphological phenotype level via MV. The dataset after selection resulted in 1854 feature vectors, 1 × 9 each, for both binary and multiclass classification. The selected features for each morphological phenotype and classification task are reported in the Supplementary Information (Table S2 and Table S3).

### 3.2 Classification results

Before performing the two classification tasks, we conducted a Principal Component Analysis (PCA) [46] to explore whether the extracted features could naturally cluster the samples according to their severity levels in a 3D projection, without the aid of machine learning. As shown in Fig. 3, in the binary classification scenario, certain phenotypes show a clearer separation between low and high severity groups, while for others the clusters overlap more. In contrast, in the multiclass setting, samples with close severity levels tend to overlap significantly, making natural clustering more challenging. In particular, in the binary classification case (top panel Fig. 3), samples tend to cluster into two main groups, with individuals belonging to classes 0/1 (red) and 2/3 (blue). The separation is evident in phenotypes B, D and F, where the overlap between the two severity clusters is negligible. Phenotypes A and C show a partial clustering, although with some mixing at the boundaries, while E and G present weaker separation.

**Figure 3.**
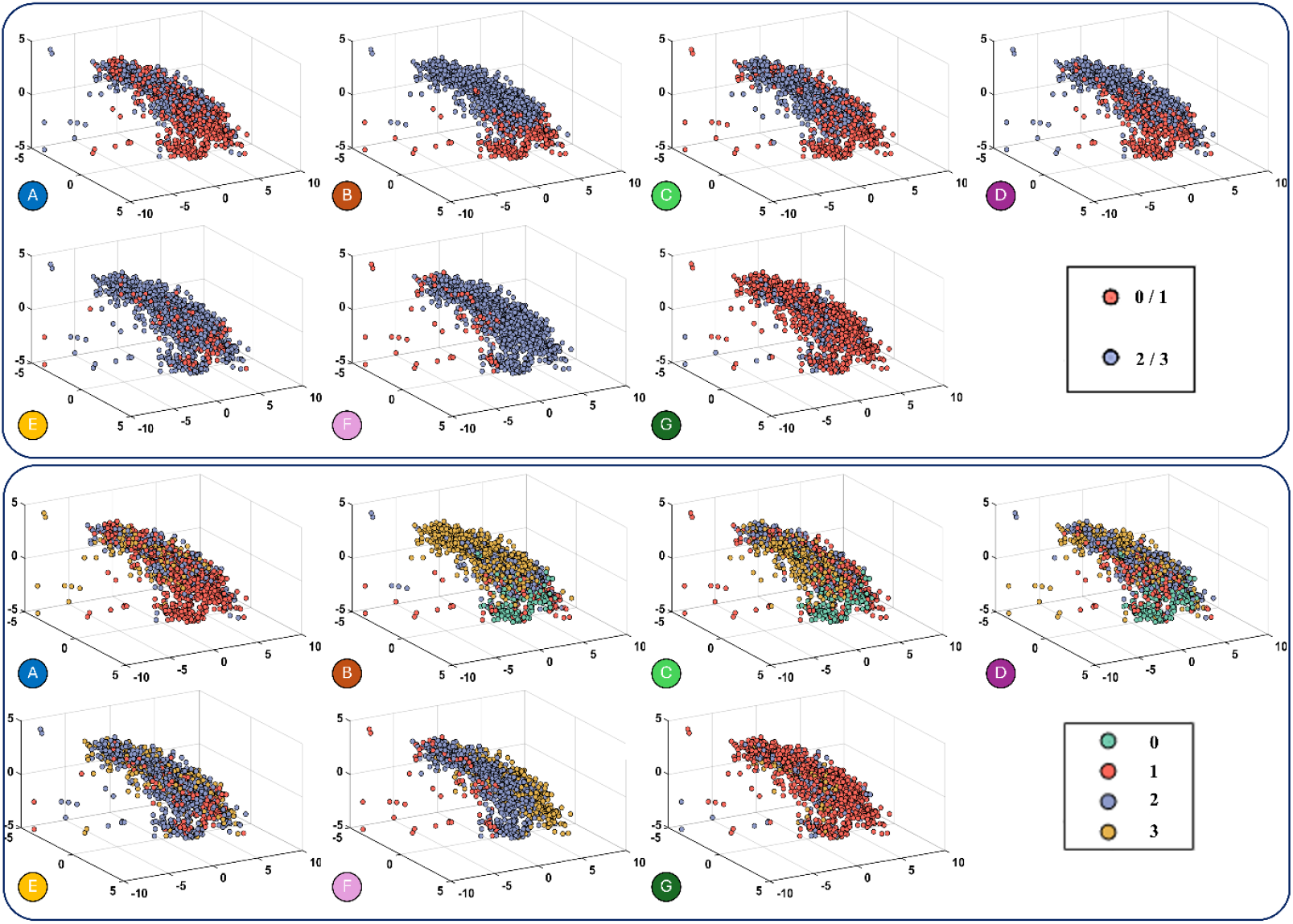
Three-dimensional Principal Component Analysis (3D PCA) plots showing the distribution of each morphological phenotype in terms of its expression severity. The top panel illustrates the binary classification task (0/1 vs 2/3), while the bottom panel displays the four classes identification task (0, 1, 2, 3). Colours indicate severity classes as specified in the legends.

In the four-class identification task, features cluster less, as expected. While the most extreme groups (0 and 3) show a soft separation, the intermediate categories (1 and 2) overlap extensively, blurring the boundaries between classes. These observations highlight the intrinsic difficulty of the problem, especially when attempting to distinguish between adjacent severity levels, and justify the use of ML approaches to effectively address the classification task. Table 1 reports the results of the binary classification task for the full set of features (37), showing the validation accuracy, test accuracy, and max voting accuracy for each phenotype.

**Table 1.**
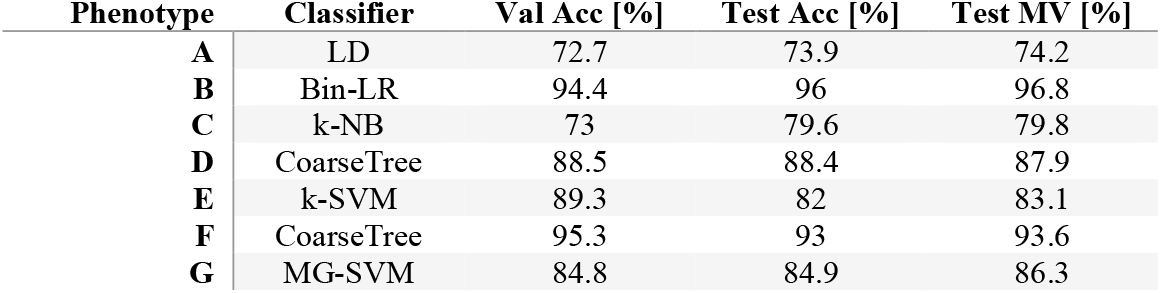
Classification results for the binary task by using 37 features. Validation, test and max voting accuracies are reported. LD: linear discriminant; Bin-LR: binary logistic regression; k-NB: kernel Naïve Bayes; k-SVM: kernel support vector machine; MG-SVM: mean gaussian support vector machine.

Table 2 reports the binary classification results obtained by using the subset selected via feature selection (9). The use of max voting across the three slices per cell led to a slight improvement in classification performance. On average, test accuracies exceed 80%, with a peak performance of 96% achieved for phenotype B. In fact, as we expected from PCA plots, phenotypes B, D, F and G have the highest accuracies, while A, C and E the lowest ones. Phenotype A remained below 75% accuracy in the binary setting and after using the selected features, the accuracy increased to exactly 75%. Notably, phenotype A is associated with alterations in nuclear morphology. The automatically selected features, listed in Table S2 of the Supplementary Information, are primarily related to descriptors of nuclear shape and structure. This suggests that tailored feature selection may help to partially compensate for the complexity in the identification of the expression severity for this phenotype. Table 3 presents the results of the multiclass identification task by using 37 features, while Table 4 by using 9 selected features. For phenotypes B, F, and G, the test accuracies reached approximately 80%, which can be considered an important outcome given the complexity of the task. In contrast, for phenotypes A, C, D, and E, accuracies fall under 70%, which aligns with expectations due to the difficulty of distinguishing between adjacent severity levels. The best results were obtained when using the full set of features, emphasizing the importance of a rich and diverse feature space for handling fine-grained severity classification.

**Table 2.** Classification results for the multiclass task by using 9 selected features. Validation, test and max voting accuracies are reported. Q-SVM: quadratic support vector machine; C-SVM: cubic support vector machine; C-KNN: coarse k-nearest neighbours; LD: linear discriminant; k-NB: kernel naïve bayes; Cos-KNN: cosine k-nearest neighbours.

| Phenotype | Classifier | Val Acc [%] | Test Acc [%] | Test MV [%] |
| --- | --- | --- | --- | --- |
| A | Q-SVM | 73.1 | 74.7 | 75 |
| B | C-SVM | 93.8 | 95.4 | 96 |
| C | C-KNN | 73.3 | 78.8 | 79 |
| D | LD | 86.5 | 89.5 | 90.3 |
| E | CoarseTree | 87.3 | 82.3 | 82.3 |
| F | k-NB | 93.6 | 93.3 | 93.6 |
| G | Cos-KNN | 76.7 | 79.8 | 81.5 |

**Table 3.** Classification results for the binary task by using 37 features. Validation, test and max voting accuracies are reported. LD: linear discriminant; EBoostT: ensemble boosted trees; EBagT: ensemble bagged trees; ESubDis: ensemble subspace discriminant.

| Phenotype | Classifier | Val Acc [%] | Test Acc [%] | Test MV [%] |
| --- | --- | --- | --- | --- |
| A | LD | 62.9 | 65.1 | 70.1 |
| B | EBoostT | 75.2 | 78.5 | 79.8 |
| C | EBagT | 65.7 | 54.8 | 56.0 |
| D | ESubDis | 63.1 | 65.3 | 65.3 |
| E | MediumTree | 66.7 | 68.8 | 71.0 |
| F | LD | 81.6 | 76.9 | 79.0 |
| G | EBagT | 80.8 | 80.9 | 81.5 |

**Table 4.**
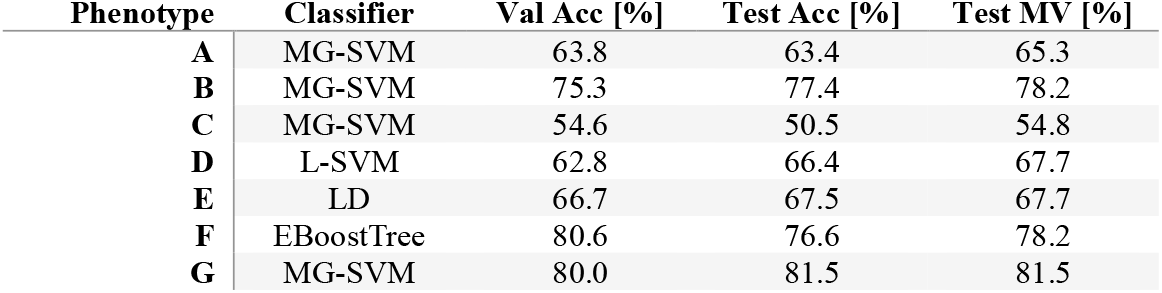
Classification results for the multiclass task by using 9 selected features. Validation, test and max voting accuracies are reported. MG-SVM: mean gaussian support vector machine; L-SVM: linear support vector machine; LD: linear discriminant; EBoostT: ensemble boosted trees.

Table 5 reports a comparison between the best result achieved using the full set of 37 features vs. the result achievable by using only the 22 features calculated from the maps of the three orthogonal planes, i.e. without considering the MIPs. It is apparent that the use of features extracted from the MIP maps brings an improvement in terms of classification accuracy despite the partial correlation existing between the information from a slice and the corresponding MIP.

**Table 5.** Classification results for the multiclass task. A comparison is reported between the full set of 37 features and the set of 22 features extracted without considering the MIP maps.

| Phenotype | Test Acc (full set) [%] | Test Acc (without MIP maps) [%] | Test MV (full set) [%] | Test MV (without MIP maps) [%] |
| --- | --- | --- | --- | --- |
| A | 65.1 | 62.4 | 70.1 | 65.3 |
| B | 78.5 | 74.2 | 79.8 | 75.0 |
| C | 54.8 | 51.9 | 56.0 | 55.7 |
| D | 65.3 | 62.9 | 65.3 | 66.9 |
| E | 68.8 | 66.4 | 71.0 | 64.5 |
| F | 76.9 | 76.6 | 79.0 | 79.0 |
| G | 80.9 | 82.3 | 81.5 | 80.7 |

### 3.3 Phenotype and severity sequence prediction

As mentioned above, biologists from our group annotated each cell by assigning a sequence of morphological phenotypes based on the analysis of the full 3D RI tomograms with added nucleus specificity. For example, a WT cell may be labelled with the following sequence: A1.B0.C0.D0.E2.F2.G1., where the numbers indicate the expression severity level of each phenotype in that particular cell. The goal of our work is to predict this entire phenotype-severity sequence for each cell. Hence, along with the assessment of the classification performance in the identification of the single phenotype expression severities, it is of interest to determine how many coexisting morphological phenotypes and associated severities are correctly predicted as a whole. Therefore, using the results from the multiclass classification task, we evaluated how many test cells had their full sequence correctly predicted by aggregating the outputs from the individual phenotype classifiers. In Fig. 4, we show the workflow of the classification pipeline for the prediction of the cell 7-digits codes. In particular, we report an example of the main auxiliary maps obtained from the same cell from which we extract the features. These features were processed independently by eight classifiers (A–G), each trained to assign a degree of severity score to each phenotype.

**Figure 4.**
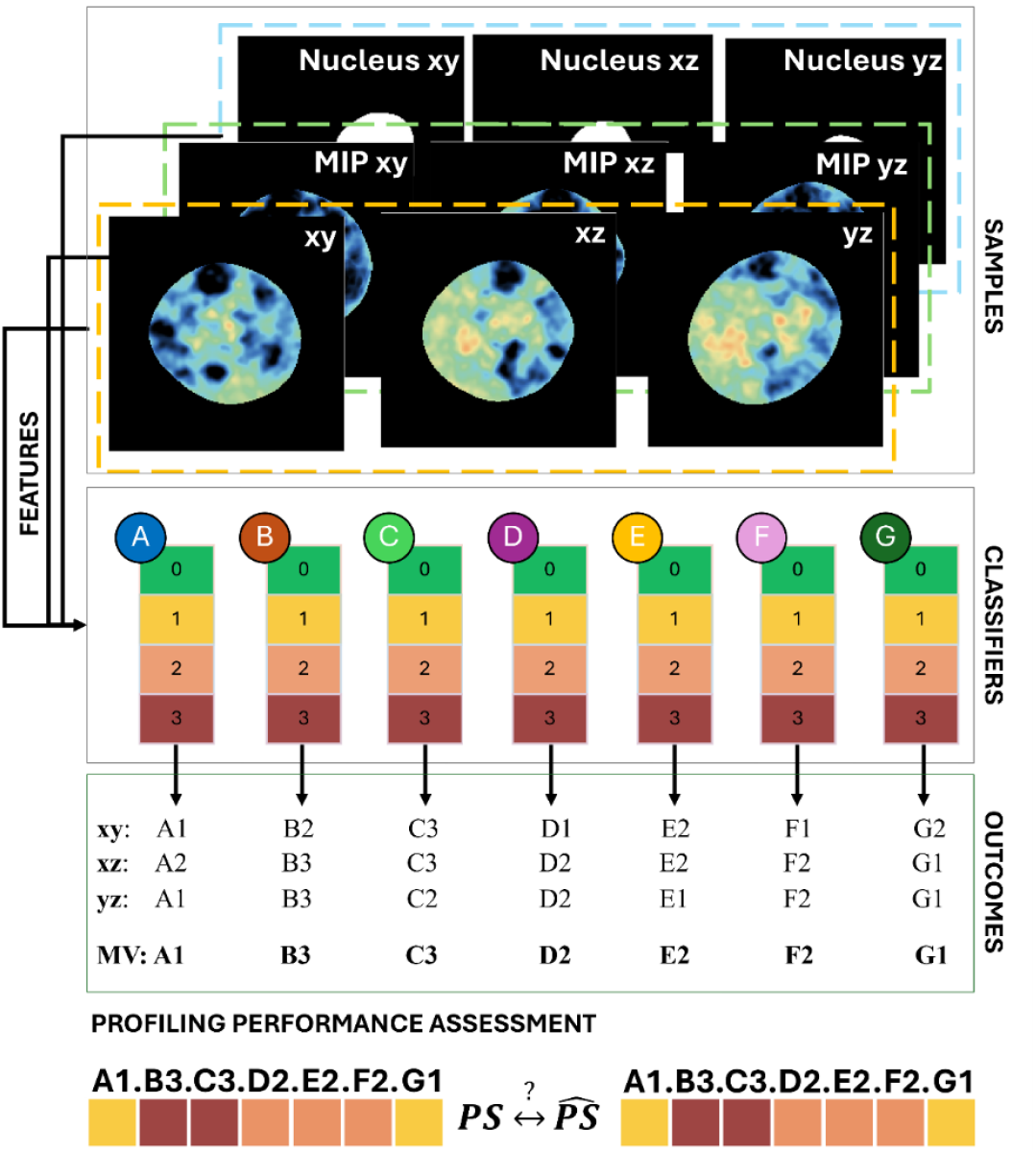
Classification framework for phenotype sequencing prediction. For each cell, the auxiliary main images considered for features extraction are the three orthogonal planes (xy, xz, yz) of RI, MIP and the nucleus support. These features are subsequently fed into multiclass classifiers (A–G). Each classifier outputs a severity score ranging from 0 to 3. The predictions from the three planes are combined, and a max voting strategy is applied to yield the final outcome for each classifier (in red). The resulting set of classifier outcomes (A1, B3, C3, D2, E2, F2, G1) is then represented as a 7-digits code to facilitate comparison between predicted phenotype sequence 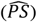 and reference phenotype sequence (PS).

For each classifier, the predictions from the three orthogonal planes were combined by MV to determine the final assigned class (bold black font in Fig. 4). For instance, classifier B produced outputs (“B2”, “B3”, “B3”) across the three orthogonal planes, leading to a final decision for “B3”. The same procedure was applied to all classifiers independently, thus increasing the final test accuracy and yielding one aggregated outcome per classifier.

The collection of final outcomes (e.g., A1, B3, C3, D2, E2, F2, G1) was then represented in a 7-digit sequence format, providing a compact visualization of the compresence of multiple morphological phenotypes that characterize in compact form the single cell and its status. This representation allowed a direct comparison between the predicted phenotype sequence 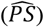 and the label phenotype sequence (PS) for each subject. Figure 5 A) shows the accuracy distribution for complete phenotypic sequence predictions for the cells belonging to the test dataset. For a considerable number of cells we achieved accuracies above 70%, with 12 cells reaching 100% accuracy in the 7-digit code prediction, meaning the full morphological sequence considered here was correctly reconstructed. The majority of predictions fall within the 60–90% range, whereas only a small number of cases remained below 40%. Figure 5 B) provides a complementary evaluation, where accuracy is computed across the total number of predicted elements. The average accuracy across samples (in this case the entire sequences) was 70.6% (median 71.4%), with a standard deviation of 18.2%. This analysis demonstrates that while achieving perfect reconstruction of the entire phenotype sequencing can be challenging, the model still maintains a solid phenotype severity prediction. In Fig. S4 of the Supplementary Information, we present a heatmap displaying the 12 test cells for which the full phenotype-severity sequence was correctly predicted, allowing for a quick visual interpretation of the phenotypic composition across these cells.

**Figure 5.**
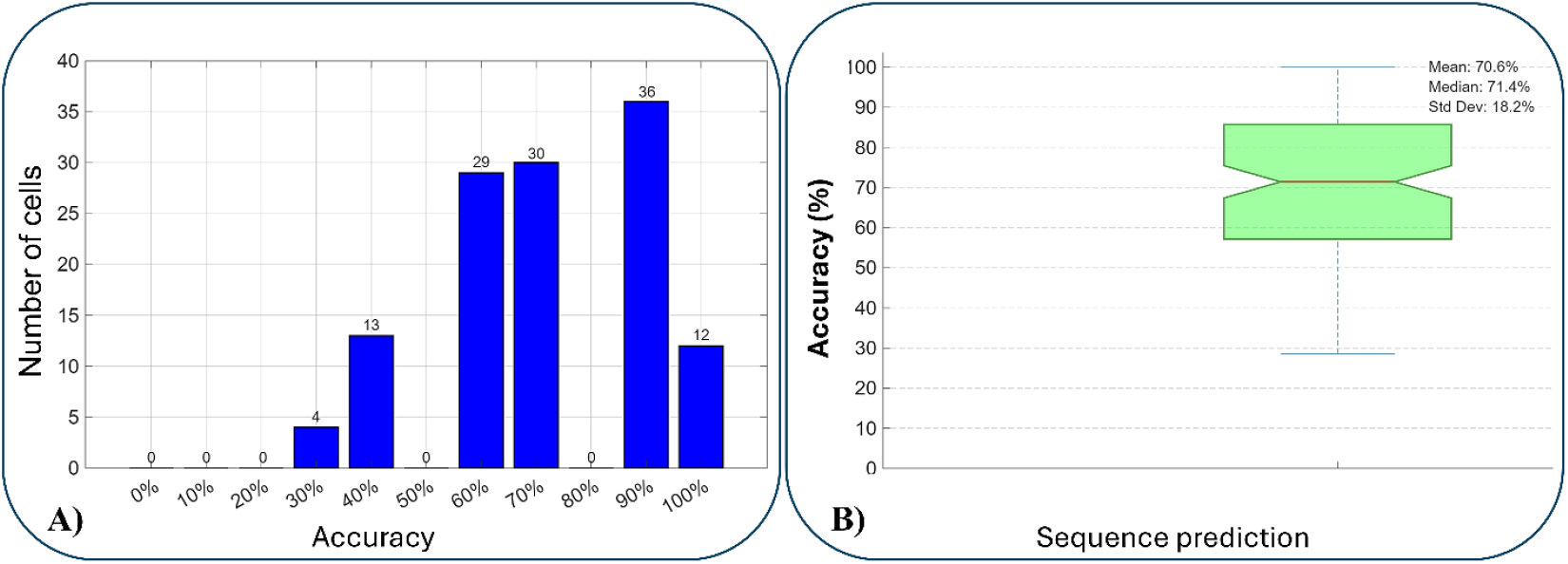
Prediction accuracy of phenotype sequences. A) Histogram showing the distribution of accuracies when the entire phenotype sequence is predicted correctly with respect to the number of test cells. Most cells reach accuracies between 60% and 90%, with a subset of 12 cells achieving 100% full-sequence prediction. B) Boxplot summarizing the overall sequence prediction performance. The mean accuracy is 70.6%, with a median of 71.4% and a standard deviation of 18.2%, indicating moderate variability across samples.

Figure 6 shows an isolevel visualization of some 3D RI tomograms obtained from the different experimental conditions under test. For each of them, we highlight the nucleus and the vacuolar compartments. We selected examples from the test set distribution of Fig. 5, showing cases of fully correct sequence prediction as well as cases for which the sequence was mostly misclassified (e.g. 40% Accuracy). For each cell, the top sequence is the ground-truth label, the bottom sequence is the prediction. For each phenotype, red and green colours mark the wrong or correct prediction, respectively. In Figure 6 we also report the corresponding morphological profiling using spider-diagrams with color-coded severity levels (see also the example in Fig. 1 D). The diagram offers a clear view of the way the different morphological phenotypes coexist within each cell and to which extent each of them is expressed.

**Figure 6.**
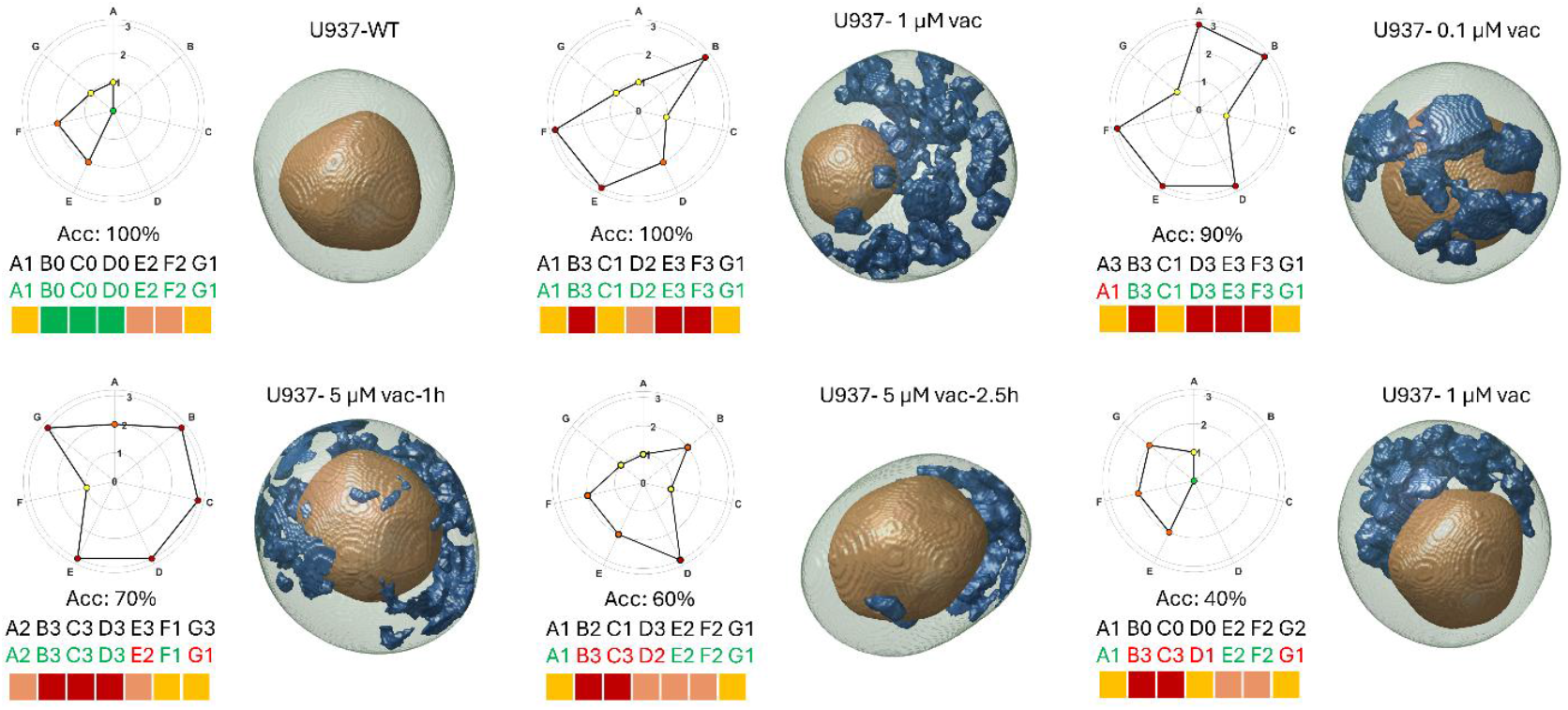
Examples of 3D RI tomograms of vacuolated cells and the corresponding estimated pattern of coexisting morphological phenotypes. vac: vacuoline. Acc: Accuracy in the overall sequence prediction. Top sequence: ground truth. Bottom, coloured sequence: estimated profile. Red/green colour in the sequence mark the wrong/correct prediction of a phenotype. Spider graphs summarize for each cell the distribution of its morphological phenotypes and their expression severity.

## 4. Discussion and conclusions

The results of our study underscore both the potential and the challenges of using label-free, 3D RI-based imaging and ML for the classification of co-occurrent morphological phenotypes and the relative degree of expression severity. The performance achieved across both binary and multiclass classification tasks highlights the discriminative power of morphometric RI-based and fractal features extracted from the three orthogonal planes of volumetric data. The variability observed between phenotypes and severity levels reflects the biological and analytical complexity of this problem. A cell is a complex system and multiple phenotypes coexist and influence each other’s morphometry. This work is a first proof of concept, carried out for the case of vacuolated mammalian cells, of the capability of HTFC in describing this compresence. In most cases, the presence or absence of a morphological phenotype in a cell is correctly assessed (binary classification). Notably, the accuracy in classifying phenotypes such as B, F, and G suggests that certain vacuoles morphologies are more pronounced, structured, or consistently captured by the defined feature vectors. Conversely, phenotypes like A, C and E are harder to be classified in terms of expression severity. These differences highlight the importance of phenotype-specific descriptors tailored to the morphological nature of each class. The challenge in multiclass classification, especially when separating close severity levels, stems from the fact that cellular changes are continuous and do not always fit well into fixed severity grades. This limitation underscores the potential use of developing classification models that can better capture graded transitions. Nevertheless, the fact that a reduced subset of features could still support robust classification in most phenotypes is encouraging, especially in real-world applications where computational efficiency and model interpretability are critical.

An important outcome of this work is the multilabel prediction of phenotype-severity sequences for individual cells. This task reflects a more realistic biological scenario, where cells often exhibit mixed or overlapping traits rather than a single phenotype. The ability to correctly reconstruct complex phenotypic compositions from imaging data represents an important step toward a more comprehensive single-cell characterization. Our findings support the use of fractal descriptors as valuable tools for quantifying this morphological complexity and in the next future differentiating between disease states. As for the benchmark case of vacuolated cells, this study could offer the basis for future assays aimed at understanding how vacuolization spreads across intracellular domains or be associated with other pathological markers, and to help in the future the diagnosis of diseases associated with the vacuoles dysfunctions, such as the family of LSDs. A data dimensionality reduction approach through the use of tomogram slices and MIP maps has been proposed in this work. This is particularly important for the calculation of the multi-scale fractal features (the computational burden is much more affordable by using 2D maps instead of the full 3D tomogram) and will be pivotal in the future to scale-up the analysis throughput of HTFC systems without losing in terms of phenotype profiling performance. On a general basis, we have also found a good agreement between the morphological phenotypes identified in the cells under test and the subpopulation they belonged, i.e. the different treatments each cell has undergone. For instance, cells treated with the largest concentration of vacuoline have been found to express high severity levels of the co-occurent phenotypes associated with the formation of cytoplasmic vacuoles. Differently, these were absent or weakly expressed for the wild type group of cells.

Despite these encouraging results, some limitations have to be pointed out. For instance, the dataset was based on cells engineering to induce vacuolization in U937 monocyte cells. Moving to diagnostic applications will require a consistent number of patient-derived samples to validate its generalizability. In conclusion, the use of orthogonal slicing, the integration of fractal features and ML approach to quantify vacuolization and classify morphological phenotypes and their expression severity within single cells introduced a novel deepening in QPI single cell analysis, where the single coexisting morphological phenotypes rather than the single cells are classified, and thus the cell is returned with its own unique 7-digits profiled sequence.

## CRediT authorship contribution statement

Marika Valentino: Methodology, Software, Writing - Original Draft; Giusy Giugliano: Investigation, Data Curation; Daniele Pirone: Software, Visualization; Fabrizio Licitra: Resources; Fulvia Vitale: Resources; Pasquale Memmolo: Formal analysis, Data Curation, Software; Lisa Miccio: Methodology, Investigation, Validation, Writing - Review & Editing; Massimo D’Agostino: Conceptualization, Resources, Validation, Funding acquisition; Pietro Ferraro: Conceptualization, Formal analysis, Writing - Review & Editing; Vittorio Bianco: Conceptualization, Supervision, Funding acquisition, Project administration.

## Data availability

Data will be made available upon reasonable request to the authors. Tomographic video sequences associated with this manuscript are available on Figshare at the following links: https://doi.org/10.6084/m9.figshare.31298062; https://doi.org/10.6084/m9.figshare.31298017; https://doi.org/10.6084/m9.figshare.31297939; https://doi.org/10.6084/m9.figshare.31297261; https://doi.org/10.6084/m9.figshare.31297858.

## Acknowledgments

This work is supported by the Prin 2022 PNRR – “Label-free cytoplasmic vacUoles pheNotyping plAykit” (LUNA) Prot. 960, 30th June 2023 – funded by the Italian Ministry of University & Research in the framework of the European Union program Next Generation EU (Project CUP: B53D23002490006).

